# Architectural chromatin interactions provide a framework for context-dependent gene regulation

**DOI:** 10.64898/2026.07.17.739237

**Authors:** Mary Likhite, Gregory Andrews, Mingshi Gao, Vivekanandan Ramalingam, Vivian Hecht, Anshul Kundaje, Jill E. Moore

## Abstract

Gene regulation depends on coordinated interactions between promoters and distal cis-regulatory elements, yet understanding how these regulatory elements communicate remains a fundamental challenge in mammalian genomics. Chromatin interaction assays provide one approach for identifying potential regulatory relationships, but interpreting the biological significance of individual interactions remains difficult; chromatin interactions comprise multiple biologically distinct classes that are only partially captured by any single assay. Here, we integrate Hi-C, RNAPII ChIA-PET, and CTCF ChIA-PET with the ENCODE Registry of candidate cis-regulatory elements (cCREs) and complementary functional genomic datasets to develop an integrative framework for classifying and interpreting promoter-centric chromatin interactions.

Using this framework, we identify a distinct class of candidate architectural promoter-enhancer interactions that are characterized by increased recurrence across cellular contexts, broader promoter connectivity, and reduced dependence on linear genomic proximity. We further show that many regulatory elements anchoring these interactions transition between enhancer and CTCF-only states while maintaining stable chromatin interactions. These dual-state regulatory elements also acquire context-specific transcription factor inputs within evolutionarily conserved architectural scaffolds, suggesting that stable chromatin architecture can be repeatedly repurposed for new regulatory functions. Genes connected to these dual-state regulatory elements are enriched for developmental and signaling pathways and exhibit increased expression specificity across cell types, consistent with specialized roles in context-dependent gene regulation. Together, our findings provide a biologically informed framework for classifying and interpreting chromatin interactions and support a model in which conserved chromatin architecture provides a stable foundation upon which new regulatory programs evolve.

## Introduction

Mammalian transcription is controlled through coordinated interactions between promoters and distal cis-regulatory elements, such as enhancers and insulators. Understanding how these elements function together to regulate gene expression is central to understanding human development, cellular identity, and disease. Although recent large-scale functional genomics efforts have dramatically expanded catalogs of regulatory elements—including our own work identifying more than 2.4 million candidate cis-regulatory elements (cCREs, (Moore et al. 2026)—the mechanisms by which these elements communicate and coordinate gene regulation remain incompletely understood.

One approach toward addressing this question is to determine which regulatory elements interact with one another in three-dimensional space, thereby identifying putative regulatory relationships. Recent advances in chromosome conformation capture technologies now enable these physical interactions to be mapped on a genome-wide scale across diverse cellular contexts (Rowley and Corces 2018). Assays such as Hi-C (Lieberman-Aiden et al. 2009; Rao et al. 2014) provide untargeted surveys of chromatin interactions, with newer protocols (The ENCODE Project Consortium 2026) enabling the identification of tens of thousands of interacting loops at approximately kilobase resolution. Targeted approaches such as ChIA-PET (Fullwood et al. 2009) enrich for interactions associated with specific regulatory proteins, including RNAPII (Li et al. 2012), the primary polymerase involved in mRNA transcription, and CTCF (Tang et al. 2015), a sequence-specific DNA-binding protein involved in chromatin organization. Together, these complementary technologies have generated rich maps of three-dimensional genome organization across diverse biological contexts.

Despite these advances, interpreting the functional significance of chromatin interactions remains challenging. Chromatin interactions are not a single biological entity but instead represent multiple modes of regulatory organization. For example, “transcriptional hubs” connect actively transcribing promoters to one another and to enhancers (Uyehara and Apostolou 2023) whereas cohesin-mediated architectural interactions organize higher-order chromatin structures (Rowley and Corces 2018). Moreover, each chromatin interaction assay captures different aspects of the interaction landscape by design, making it difficult to interpret regulatory element connectivity using any single assay in isolation. These challenges motivate integrative approaches that combine chromatin interaction maps with complementary functional genomic annotations to distinguish biologically distinct categories of chromatin interactions.

Here, we integrate three dimensional chromatin interaction assays with cCREs and their underlying biochemical signals, transcriptional measurements, and sequence-based regulatory models across six deeply profiled human cell lines to develop an integrative framework for interpreting promoter-centric chromatin interactions. Using this framework, we identify distinct classes of promoter interactions with different organizational and regulatory properties. We further identify a class of architectural regulatory elements that transition between enhancer and CTCF-only states across cellular contexts while maintaining stable chromatin interactions and characterize the sequence and evolutionary features associated with these transitions. Finally, we show that these elements preferentially connect genes involved in developmental and context-dependent biological programs, supporting a model in which stable chromatin architecture provides a framework for context-dependent gene regulation.

## Results

### Integrating complementary dimensions of regulatory organization

To characterize the biological context of three-dimensional chromatin interactions, we curated complementary datasets generated by the ENCODE Consortium (The ENCODE Project Consortium 2026) across six deeply profiled human cell lines spanning diverse developmental origins and cellular states: Caco-2, HCT116, HepG2, K562, MCF 10A, and MCF-7 (**Fig. 1A**). For each cell line, we integrated cCRE annotations together with their underlying biochemical signals (DNase-seq, histone modifications, and CTCF ChIP-seq), sequence-level regulatory predictions from ChromBPNet (Pampari et al. 2025), steady-state and nascent transcription measurements from RNA-seq and Bru-seq, and three-dimensional chromatin interactions identified by Hi-C, RNAPII ChIA-PET, and CTCF ChIA-PET (**Supplemental Table S1A**). To ensure that our conclusions reflected reproducible patterns rather than observations from any single cell line, we performed all analyses independently in each of the six cell lines and only reported trends consistently observed across cell types. Unless otherwise noted, figures highlight results from MCF-7 because this cell line exhibited characteristics representative of the six cell lines and has been extensively characterized in previous studies of three-dimensional genome organization (Li et al. 2012).

**Figure 1.**
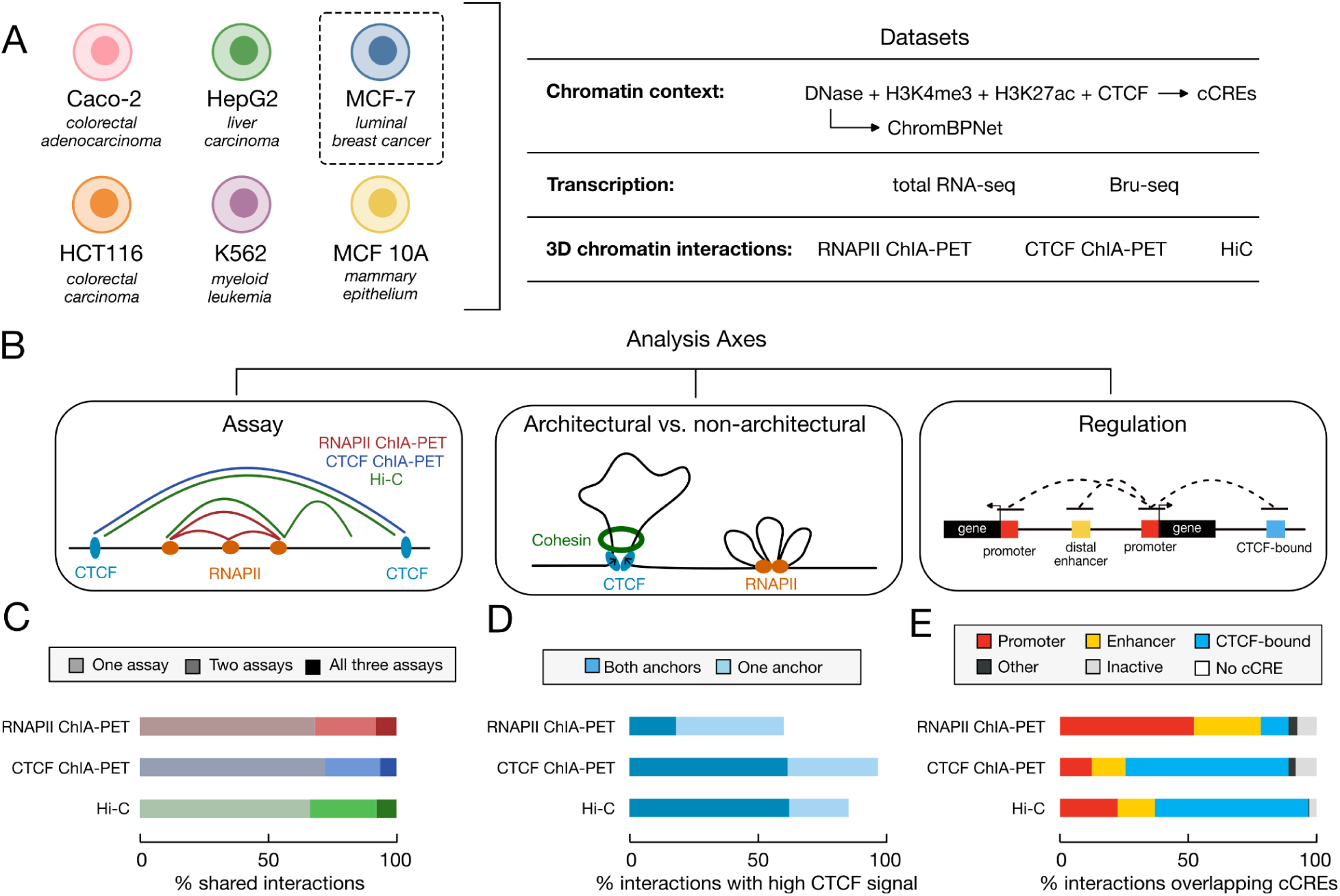
Integrative framework for interpreting promoter-centered chromatin interactions. (*A*) Overview of the six human cell lines and functional genomic datasets analyzed in this study. Chromatin context was characterized using elements from the Registry of cCREs, DNase-seq, histone modifications, CTCF ChIP-seq, and ChromBPNet attribution scores. Transcription was assessed using total RNA-seq and Bru-seq, and three-dimensional chromatin interactions were characterized using RNAPII ChIA-PET, CTCF ChIA-PET, and intact Hi-C. (B) Conceptual overview of the three complementary axes used to interpret chromatin interactions. Interactions were stratified according to (i) chromatin interaction assay, (ii) evidence for architectural organization, and (iii) the regulatory classes of interacting cCREs. (*C*) Bars plots depicting the overlap of chromatin interactions identified by RNAPII ChIA-PET, CTCF ChIA-PET, and intact Hi-C. Bars indicate the percentage of interactions detected by a single assay (light), two assays (medium), or all three assays (dark). (*D*) Barplots depicting the fraction of interactions in each assay overlapping high signal CTCF-bound cCREs at one (light blue) or both (dark blue) interaction anchors in MCF-7. Corresponding values for all other cell lines are provided in **Supplemental Table S1F.** Interactions overlapping high signal CTCF-bound cCREs at both anchors were designated as candidate architectural interactions. (*E*) Distribution of cCRE classes overlapping interaction anchors for each chromatin interaction assay in MCF-7. Corresponding values for all other cell lines are provided in **Supplemental Table S1G.**

Across the six cell lines, we analyzed approximately 917,630 chromatin interactions whose anchors overlap 315,474 unique cCREs active in at least one of the six cell types (**Supplemental Table S1B, Supplemental Dataset S1**). Despite the scale of this resource, the total number of detected interactions varied substantially across datasets, particularly for ChIA-PET experiments. RNAPII and CTCF ChIA-PET datasets ranged from 14 to 85 thousand interactions per cell type, whereas Hi-C datasets were somewhat more consistent, ranging from 44 to 80 thousand interactions. These differences reflect substantial technical variability between datasets, complicating direct comparisons based on interaction counts alone. Therefore to enable systematic comparisons across assays and cell lines, we developed a framework that evaluated interactions along three complementary axes: (1) the chromatin interaction assay used to detect the interaction, (2) evidence for architectural organization, and (3) the regulatory classes of interacting cCREs (**Fig. 1B**).

We first assessed the extent of overlap among Hi-C, RNAPII ChIA-PET, and CTCF ChIA-PET interactions within the same cell line. Although the three assays identified overlapping interactions, each also detected substantial numbers of assay-specific interactions (average 64%, **Fig. 1C, Supplemental Table S1C**), highlighting that they capture complementary subsets of the chromatin interaction landscape. Further comparison of interaction properties, including genomic distance, cCRE composition, and CTCF support, suggested that these assay-specific interactions differed systematically in their regulatory characteristics (**Fig. 1D,E**; **Supplemental Fig. S1**). Rather than viewing these differences solely as inconsistencies between assays, we reasoned that integrating complementary chromatin interaction assays could provide additional insight into the mechanisms underlying chromatin interactions.

The second axis we considered was evidence for architectural organization. Architectural chromatin loops are a major organizing feature of the mammalian genome, helping define regulatory domains within which promoters and distal regulatory elements interact (Rowley and Corces 2018). These structures primarily arise through cohesin-mediated loop extrusion and are frequently anchored by convergently oriented CTCF binding sites. To distinguish interactions likely reflecting chromatin architecture from those associated with other forms of regulatory organization, we defined a set of *candidate architectural interactions* across all six cell types (**Supplemental Table S1D**). Specifically, interactions were designated as candidate architectural interactions if both anchors overlapped cCREs with high CTCF ChIP-seq signal. This approach provided several practical advantages, including a consistent architectural definition across all six cell lines, direct compatibility with cCRE annotations, and independence from cell type-specific cohesin ChIP-seq availability. Importantly, this architectural designation based on CTCF co-occupancy at interaction anchors was concordant with alternative approaches such as those using cohesin co-occupancy or CTCF motif orientation (**Supplemental Table S1E, Supplemental Methods**), supporting its use as a conservative proxy for architectural organization.

Stratifying interactions by assay revealed marked differences in architectural support (**Fig. 1D, Supplemental Table S1F**). On average, candidate architectural interactions comprised just over half of all interactions detected by Hi-C, but only 11% of interactions identified by RNAPII ChIA-PET. As expected, CTCF ChIA-PET interactions were strongly enriched for CTCF occupancy, with 95% containing at least one anchor overlapping a CTCF-bound cCRE. However, the fraction classified as candidate architectural interactions (both anchors with CTCF occupancy) varied substantially across cell lines (29–90%), inversely correlating with the total number of interactions per dataset (Pearson’s correlation coefficient = −0.94, *p* = 0.004). This pattern suggests that technical differences in assay sensitivity and interaction-calling depth influence the proportion of candidate architectural interactions recovered, with deeper datasets identifying proportionally more non-architectural interactions. Despite this variability, all CTCF ChIA-PET datasets remained significantly enriched for candidate architectural interactions relative to RNAPII ChIA-PET (Fisher’s exact test, p < 2.2 × 10^−16^).

The final axis we considered was the predicted function of interacting regulatory elements. We intersected interaction anchors with cell type-specific annotations from the Registry of cCREs (Moore et al. 2026). Because CTCF binding plays a central role in chromatin architecture, we extended the standard Registry classification scheme by defining a dedicated class of CTCF-bound cCREs based on strong evidence of CTCF occupancy independent of chromatin accessibility or histone modification state (**Methods**). This class differs from the standard Registry framework, which requires DNase accessibility as a prerequisite for active cCRE classification. Across all assays, more than 98% of interactions overlapped at least one cCRE active in the surveyed cell type, indicating that chromatin interactions are highly enriched for regions with regulatory potential (Moore et al. 2026).

Stratifying interactions by overlapping cCRE classes revealed notable differences among assays (**Fig. 1F**, **Supplemental Table S1G, Supplemental Methods**). Unsurprisingly, RNAPII ChIA-PET interactions were strongly enriched for promoters, with an average of 64% of interactions overlapping a promoter cCRE. In contrast, Hi-C and CTCF ChIA-PET interactions were enriched for CTCF-bound cCREs, which comprised an average of 50% and 64% of interactions, respectively. Overlapping enhancer cCREs were observed across all assays, but were most prominent in RNAPII ChIA-PET datasets (26% of interactions). Inactive and non-cCRE annotations accounted for only a small minority of interaction anchors across all assays (**Supplemental Table S1G**). These observations indicate that different assays preferentially capture distinct classes of regulatory elements, with RNAPII ChIA-PET emphasizing promoter-centric regulatory interactions and Hi-C and CTCF ChIA-PET exhibiting greater enrichment for interactions associated with chromatin architecture. Altogether, these findings demonstrate that individual chromatin interaction assays capture distinct and overlapping subsets of regulatory organization, motivating an integrated framework that jointly considers assay type, architectural support, and regulatory element function.

### Architectural promoter interactions represent a distinct class of promoter-centered connectivity

Because our goal was to understand how three-dimensional genome organization contributes to transcriptional regulation, we focused subsequent analyses on promoter-centric interactions. On average, 84% of active promoters in each cell line participate in at least one promoter-centric interaction (**Supplemental Table S2A**). Drawing inspiration from earlier studies (Sanyal et al. 2012), we classified promoter-centric interactions into three categories (**Supplemental Table S1D, Supplemental Datasets S2 & S3, Supplemental Methods**): interactions between two promoters, (P–P); interactions linking promoters to enhancers (P–E); and interactions connecting promoters to CTCF-bound cCREs (P–C). To generate non-overlapping interaction classes, cCRE annotations were assigned hierarchically (promoter > enhancer > CTCF). Across the six cell lines, these three categories accounted for an average of 76% of all promoter-centric interactions, with the remainder comprising other cCREs classes or inactive cCREs (**Supplemental Table S2A**). Among the three major categories, P–C interactions were the most abundant, comprising 29% of promoter-centric interactions on average, followed by P–E (26%) and P–P (21%) interactions. (**Fig. 2A, Supplemental Table S2A**).

**Figure 2.**
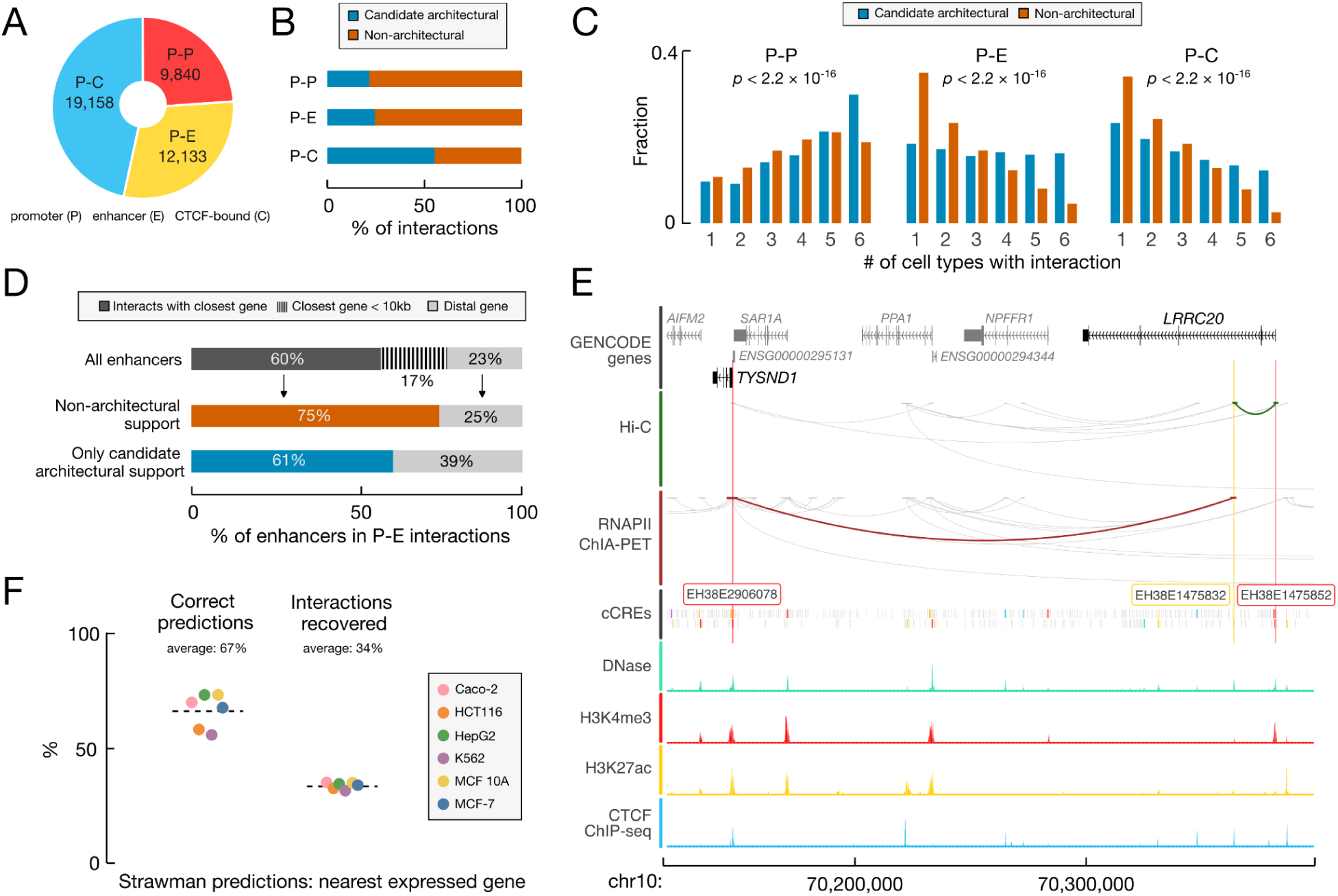
Properties of promoter-centric interactions. (*A*) Pie chart depicting the percentage of promoter-centric interactions classified as promoter-promoter (P-P), promoter-enhancer (P-E), promoter-CTCF-bound (P-C). Values shown are from MCF-7; corresponding values for all cell lines are provided in **Supplemental Table S2A**. (*B*) Bar plots depicting the fraction of each promoter-centric interaction class classified as candidate architectural (blue) or non-architectural (orange) in MCF-7. Corresponding values for all other cell lines are provided in **Supplemental Table S2C.** (*C*) Bar plots depicting the recurrence of promoter-centric interactions in MCF-7 across the six cell lines stratified by architectural support (as defined in *B*). P-values correspond to Wilcoxon rank-sum tests. Corresponding values for all other cell lines are provided in **Supplemental Table S2D.** (*D*) Bar plots depicting the target-gene assignment for enhancers participating in P-E interactions in MCF-7 cells. Shown are the fractions of enhancers interacting with their nearest promoter (gray, orange, blue), located within 10 kb of the nearest TSS (below the effective resolution of current chromatin interaction assays; dashed lines), or interacting exclusively with more distal promoters (light gray). Below barplots are stratified by enhancers based on their interaction classes: those with non-architectural interactions (orange) and those with only candidate architectural interactions (blue). Enhancers located within 10 kb of the nearest gene are removed in these comparisons. Corresponding values for all other cell lines are provided in **Supplemental Table S2F.** (*E*) Genome browser tracks from MCF-7 cells at the *TYSND1*-*LRRC20* locus. Enhancer cCRE EH38E1475832 (yellow highlight) is connected to the promoter (EH38E1475852, red highlight) of the closest gene *LRRC20* by an non-architectural interaction detected by Hi-C (green arc). EH38E1475832 is also connected to the promoter (EH38E2906078, red highlight) of distal gene *TYSND1* by a candidate architectural interaction detected by RNAPII ChIA-PET (red arc). Bottom tracks include are MCF-7 cCREs are colored by class with supporting biochemical signals DNase-seq (green), H3K4me3 ChIP-seq (red), H3K27ac ChIP-seq (yellow), CTCF ChIP-seq (blue). (*F*) Performance of a nearest-expressed-gene method for predicting enhancer-promoter relationships across the six cell lines. While the nearest expressed gene correctly identified the interacting promoter for an average of 66% of enhancers, it recovered only a median of 34% of all P-E interactions.

P–P, P–E, and P–C interactions were detected across all three chromatin interaction assays, although the relative contribution of each assay differed among the categories (**Supplemental Table S2B**). P-P and P-E interactions were supported primarily by RNAPII ChIA-PET and Hi-C, which together accounted for an average of 91% of P-P interactions and 86% of P-E interactions. P–C interactions, however, received substantial support from all three assays with an average of 24% from RNAPII ChIA-PET, 32% from CTCF ChIA-PET and 44% from Hi-C. Because candidate architectural interactions were defined independently of promoter-centric interaction category, each P–P, P–E, and P–C interaction could be further designated as either candidate architectural or non-architectural. The three promoter-centric categories differed substantially in the fraction of interactions designated as candidate architectural (Fig. 2B, **Supplemental Table S2C**). The majority of P–C interactions were candidate architectural (cross-cell-line average 55%), compared with smaller fractions of P–E (∼19%) and P–P (∼20%). Additionally, consistent with previous studies of cohesin-associated looping, candidate architectural interactions spanned significantly longer genomic distances than their non-architectural counterparts, increasing an average of 22 kb for P-E and 46 kb for P-C (**Supplemental Fig. S2**). Unlike P-E and P-C interactions, architectural P-P interactions did not exhibit consistent distance differences across the six cell lines. Together, these observations indicate that promoter-centric interaction classes differ in both their regulatory composition and their association with chromatin architecture.

To assess recurrence of promoter-centric interactions across cell types, we calculated the number of cell lines in which each interaction was observed (**Fig. 2C**, **Supplemental Table S2D**). P-P interactions were the most broadly shared across the six cell lines, with an average of 58% of interactions observed in at least four of the six cell lines. In contrast, P-E and P-C interactions were substantially more cell type-specific, with only 30% and 40% observed in at least four cell lines, respectively. However, within both classes, candidate architectural interactions were more frequently detected across multiple cell types. On average, 52% of architectural P-E interactions and 48% of architectural P-C interactions were observed in at least four cell types, compared with only 25% and 31% of non-architectural interactions, respectively. Together, these observations suggest that promoter-centric interaction categories capture one dimension of chromatin interaction organization and that architectural organization provides additional biological context. We therefore examined each interaction class in greater detail.

Across the six cell lines, an average of 43% of genes with active promoters participated in at least one P-P interaction (**Supplemental Table S2E**). These genes exhibited the highest expression levels, with an average expression of 9.9 transcripts per million (TPM) across the six cell lines, compared with 6.3 TPM for genes associated exclusively with P-E and/or P-C interactions and 3.8 TPM for genes lacking promoter-centric interactions altogether (Pairwise Wilcoxon tests with FDR correction, *p* < 7.9 × 10^−11^, **Supplemental Fig. S3**). Genes connected through P-P interactions also exhibited greater expression correlation with their interacting partner than randomly selected promoter pairs (average 0.47 vs. 0.12, Wilcoxon tests, *p* < 2 × 10^−16^; **Supplemental Fig. S4**). Together, these results are consistent with previous studies suggesting that P-P interactions preferentially connect transcriptionally active genes within shared regulatory neighborhoods.

We next examined the organizational properties of P-E interactions. Across the six cell lines, an average of 44% of genes with active promoters participated in at least one P-E interaction (**Supplemental Table S2E**). After accounting for annotation-dependent biases in enhancer-gene assignment (**Supplemental Methods**), we found that approximately 60% of TSS-distal enhancers interacted with the promoter of their closest gene (**Supplemental Table S2F**. Of the remaining enhancers, 17% were located within 10 kb of the nearest promoter, a distance below the effective resolution of the 3D interaction assays (**Supplemental Fig. S5**), leaving only ∼24% of enhancers linked exclusively to more distal genes. Architectural and non-architectural interactions exhibited subtle but consistent differences in target-gene selection. After excluding enhancers within 10 kb of the nearest promoter, enhancers participating in non-architectural interactions more frequently targeted their nearest promoter (73%) than enhancers connected exclusively through candidate architectural interactions (54%; **Fig. 2E**). Enhancers participating in candidate architectural interactions also were connected to a greater number of genes on average compared to enhancers lacking candidate architectural interactions (2.9 versus 1.9 genes per enhancer, Wilcoxon test p < 2 × 10^−16^; **Supplemental Fig. S6**).

To evaluate the practical implications of these organizational differences, we implemented a simple “closest gene” assignment strategy linking each enhancer to the nearest expressed protein-coding gene (TPM > 1). This approach correctly identified an average of 66% of observed enhancer-promoter contacts but recovered only 34% of the contacts detected by chromatin interaction assays (**Fig. 2G, Supplemental Table S2F**). Thus, while proximity frequently identifies a nearby interacting promoter, it fails to capture the majority of chromatin contacts detected by three-dimensional interaction assays. This limited recovery likely reflects, in part, the tendency of some enhancers to contact multiple promoters, particularly those participating in candidate architectural interactions. This limited recall likely reflects, in part, the tendency of some enhancers to contact multiple promoters, particularly those participating in candidate architectural interactions. For example, enhancer EH38E1475832 is linked to the nearby promoter of *LRRC20* through a non-architectural interaction and to the more distal promoter of *TYSND1* through a candidate architectural interaction (**Fig. 2F**). Our closest-expressed-gene strategy correctly identifies the local *LRRC20* contact but fails to recover the more distal *TYSND1* contact. Together, these observations suggest that incorporating chromatin architecture can refine, rather than replace, proximity-based models of enhancer-promoter connectivity. Although chromatin interactions provide candidate regulatory relationships rather than direct evidence of regulatory function, these results demonstrate that long-range architectural interactions frequently connect enhancers to promoters beyond their nearest expressed gene.

We next asked whether architectural P-C interactions exhibited organizational properties similar to those observed for architectural P-E interactions. Across the six cell types, an average of 50% of genes with active promoters participated in at least one P-C interaction. As observed for P-E relationships, architectural P-C interactions were less frequently associated with the nearest gene than non-architectural P-C interactions (38% versus 54%; **Supplemental Table 2G**). Architectural CTCF-bound cCREs were also linked to a greater number of genes on average than non-architectural CTCF-bound cCREs (1.9 versus 1.6 genes, Wilcoxon test, *p* < 3.4 × 10^−4^; **Supplemental Fig. S6**). Together, these observations identify candidate architectural interactions as a distinct category of promoter-centric interactions characterized by greater recurrence across cell types, reduced dependence on linear genomic proximity, and broader promoter connectivity than non-architectural interactions.

### Dual-state regulatory elements emerge within recurrent candidate architectural interactions

To better understand how the unique properties of candidate architectural interactions impact gene regulation, we compared these interactions across the six cell lines. Surprisingly, despite being highly recurrent across cell lines (**Fig. 2C**), many architectural P-E interactions did not retain the same regulatory classification. On average, when architectural P-E interactions were observed in other cell types, 36% were classified as P-C interactions rather than retaining a P-E classification (45%, **Fig. 3A**, **Supplemental Fig. S3A**). HCT116 represented a notable exception. Across comparisons involving the other five cell lines, recurrent architectural P-E interactions were more likely to retain a P-E classification in HCT116 (59% versus 42% on average across other cell types) and less likely to be classified as P-C interactions (22% versus 39%). These observations raised the possibility that candidate architectural interactions are frequently maintained across cellular contexts while the regulatory state of their anchors changes.

**Figure 3.**
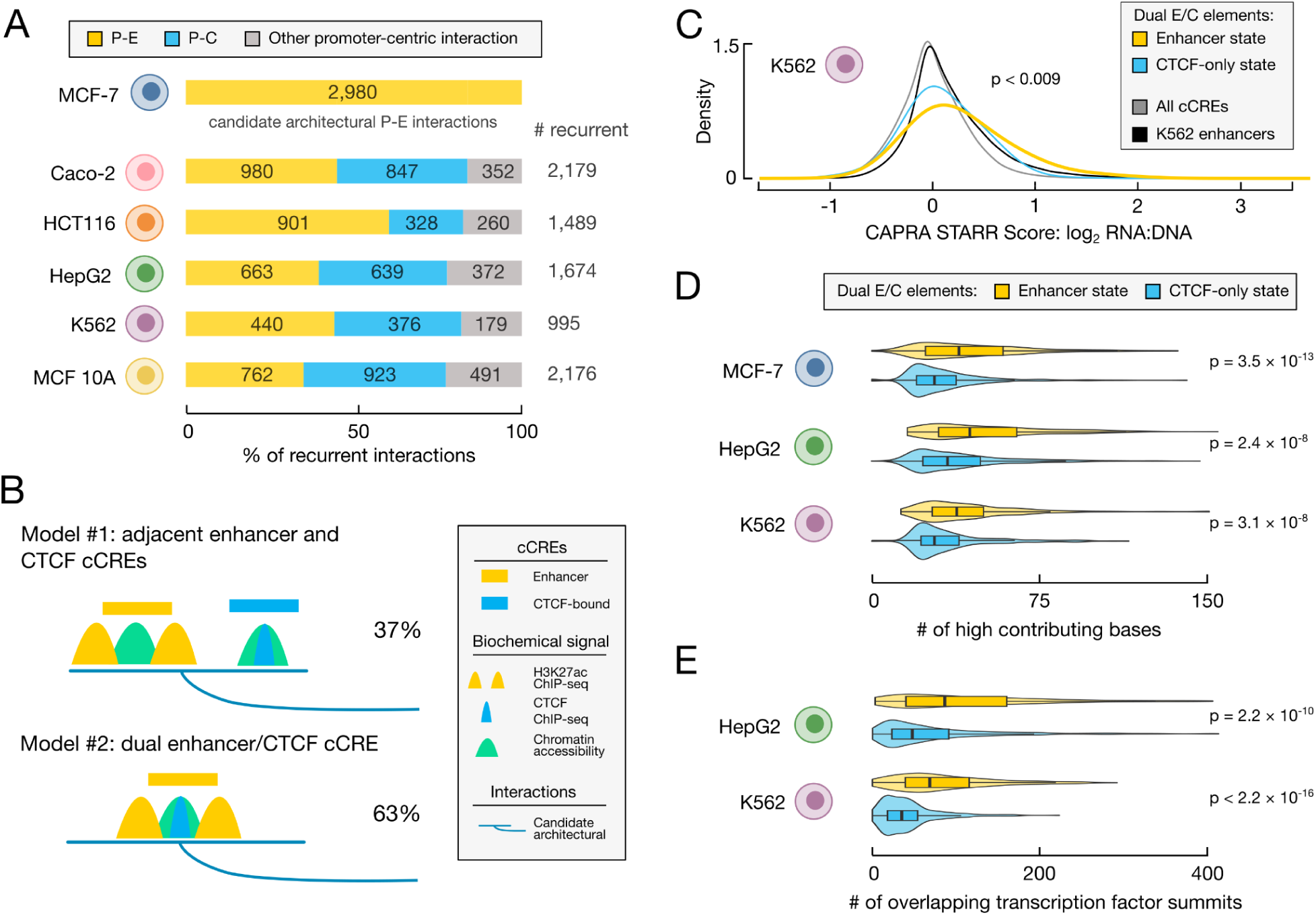
Candidate architectural interactions support regulatory state transitions within dual E/C elements. (*A*) Barplots depicting the classification of recurrent MCF-7 candidate architectural P-E interactions (N=2,980) across the other five cell lines. Percentages of interactions that are classified as P-E (yellow), P-C (blue), or other promoter-centric (gray) are shown for each cell line. The number of recurrent interactions between MCF-7 and each cell line is shown on the right. Corresponding values for all other cell line comparisons are provided in **Supplemental Table S3A**. (*B*) Schematic of two models that could explain transitions between P-E and P-C interaction classifications. In *model 1*, enhancer activity arises from a regulatory element adjacent to the CTCF-bound cCRE participating in the architectural interaction. In *model 2*, a single cCRE transitions between enhancer and CTCF-only states while maintaining participation in the same candidate architectural interaction. The majority of observed transitions (63%) were consistent with model 2. (*C*) Density plots of CAPRA STARR-seq scores for dual E/C elements in K562 cells. Dual E/C elements in the enhancer state (yellow) had higher scores than all other cCRE groups: dual E/C elements in the CTCF-only state (blue), all K562 cCREs (black) and randomly selected cCREs (gray). P-value shown corresponds to highest p-value from pairwise Wilcoxon rank sum tests with FDR correction (*D*) Violin-boxplots depicting the number of bases in dual E/C elements with high ChromBPNet attribution scores. For each of the three cell lines, dual E/C elements were stratified by enhancer state (yellow) or CTCF-only state (blue). P-values correspond to Wilcoxon rank-sum tests. (*E*) Violin-boxplots depicting overlap of transcription factor peak summits with dual E/C elements stratified by state as defined in *D.* P-values correspond to Wilcoxon rank-sum tests.

Two biological mechanisms could explain this pattern (**Fig. 3B**). First, enhancer activity could arise from a regulatory element adjacent to the CTCF-bound cCRE mediating the architectural interaction (model 1). In this scenario, separate enhancer and CTCF-bound cCREs overlap the same interaction anchor. Alternatively, the CTCF-bound cCRE itself could acquire enhancer signatures in a cell type-specific manner, resulting in a single CTCF-bound regulatory element that transitions between CTCF-only and enhancer states (model 2). To distinguish between these possibilities, we focused on candidate architectural interactions where the non-promoter anchor overlapped exactly one enhancer cCRE, allowing us to unambiguously determine whether the enhancer and CTCF-bound annotations mapped to the same underlying element (**Supplemental Fig. S3B**, **Supplemental Methods**).

In 63% of cases the enhancer and CTCF-bound annotations mapped to the same underlying cCRE (model 2). Thus, despite the potential for enhancer activity to arise at neighboring regulatory elements, regulatory state transitions most frequently occurred within a single nucleosome-free region (**Fig. 4C**). We refer to these cCREs as dual E/C elements because they can exhibit either enhancer or CTCF-only regulatory states depending on cellular context. Throughout the remainder of the manuscript, dual E/C elements exhibiting enhancer signatures are referred to as the enhancer state, whereas dual E/C elements lacking enhancer signatures are referred to as the CTCF-only state.

**Figure 4.**
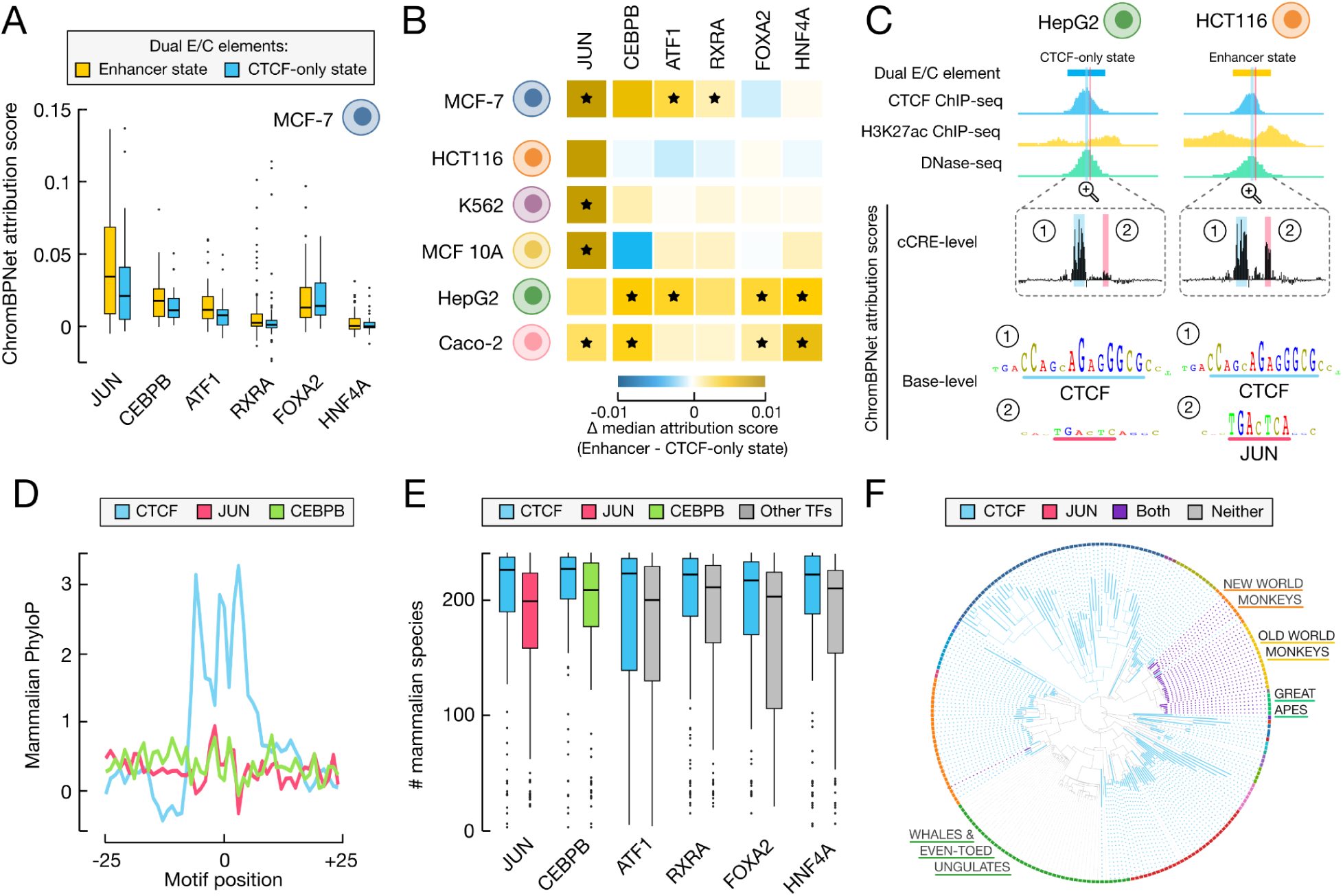
Dual E/C elements acquire context-specific regulatory inputs within evolutionarily conserved architectural scaffolds. (A) Boxplots of ChromBPNet attribution scores in MCF-7 at experimentally supported transcription factor motif instances for dual E/C elements stratified enhancer (yellow) and CTCF-only states (blue). A representative set of transcription factors are shown. Corresponding values for all other cell lines and transcription factors are provided in **Supplemental Table S4A**. (B) Heatmap depicting the median change in ChromBPNet attribution scores (enhancer state − CTCF-only state) for transcription factor motifs featured in **A** across the six cell lines. Stars indicate *p* < 0.05 by Wilcoxon rank sum test. Corresponding values for all other cell lines and transcription factors are provided in **Supplemental Table S4A**. (*C*) Genome browser tracks at dual E/C element (EH38E2226676) illustrating transition from a CTCF-only state (HepG2, left) to an enhancer state (HCT116, right). Tracks include cCREs colored by state (enhancer: yellow, CTCF: blue) with supporting biochemical signals CTCF ChIP-seq (blue), H3K27ac ChIP-seq (yellow), DNase-seq (green). Below are zoomed in views of ChromBPNet attribution scores at the cCRE (top) and base pair level (bottom) with two highlighted motifs: 1, CTCF in blue and 2, JUN in pink. This locus across all six cell lines is detailed in **Supplemental Figure S9**. (*D*) Aggregate plots depicting the average mammalian PhyloP scores across CTCF (blue), JUN (pink) and CEBPB (green) motif instances within dual E/C elements. (*E*) Boxplots depicting the number of mammalian species with conserved motif instances for CTCF vs other transcription factor motifs in the same dual E/C elements. Conserved motif instances are defined as >75% sequence similarity between the species and human. Corresponding values for all other transcription factors and corresponding p-values are summarized in **Supplemental Table S4B**. (*F*) Phylogenetic tree depicting conservation of the CTCF and JUN motif sites in representative dual E/C element EH38E2226676 shown in panel **C**. The CTCF motif is conserved across all mammalian lineages except for whales and even-toed ungulates, whereas the adjacent JUN motif is largely restricted to primates (great apes, old world monkeys and new world monkeys).

To determine whether enhancer-state dual E/C elements exhibit enhancer activity, we analyzed per-cCRE STARR-seq scores in K562 cells using our CAPRA method (Moore et al. 2026). Enhancer-state dual E/C elements exhibited substantially greater STARR-seq activity than randomly selected cCREs (median = 0.19 vs. −0.02, *p* = 7.6 × 10^−15^; **Fig. 3C**) and comparable or slightly greater activity than K562 enhancers participating in non-architectural promoter interactions (median = 0.09, *p* = 0.009). Consistent with their K562 enhancer annotation, enhancer-state dual E/C elements also exhibited significantly greater STARR-seq activity than dual E/C elements in the CTCF-only state (median = 0.06, *p* = 4.7 × 10^−4^; **Fig. 3C**). Together, these observations demonstrate that enhancer-state dual E/C elements represent active enhancers rather than annotation artifacts arising from nearby chromatin state changes. Moreover, their enhancer activity is concordant with the regulatory state assigned in K562 cells.

We next investigated how dual E/C elements transition from the CTCF-only state to the enhancer state while retaining their underlying architectural role. We hypothesized that the enhancer state arises through recruitment of additional transcription factors within the same nucleosome-free region. To investigate this possibility, we quantified the fraction of bases within dual E/C elements exhibiting high ChromBPNet attribution scores, which identify base pairs predicted to contribute strongly to local chromatin accessibility.

Across five of six cell types, enhancer-state dual E/C elements contained more high-attribution bases than their CTCF-only counterparts, with median increases ranging from 8–14 high-attribution base pairs (**Fig. 3D**, **Supplemental Fig. S7A**), suggesting that relatively modest sequence differences may underlie the regulatory state transition. HCT116 represented the sole exception, exhibiting similar attribution profiles between enhancer and CTCF-only states. Results were also consistent when attribution profiles were quantified as the fraction of high-attribution bases (**Supplemental Fig. S7B**). Consistent with this observation, enhancer-state dual E/C elements also exhibited substantially greater transcription factor occupancy (**Supplemental Dataset S4**). In HepG2, enhancer state dual E/C elements overlapped a median of 86 transcription factor ChIP-seq peak summits compared to a median of 48 for CTCF state elements (Wilcoxon test, *p* < 2.2 × 10^−10^, **Figure 3E**). We observed a similar pattern in K562 with overlaps of 68 vs 36, respectively (Wilcoxon test, *p* < 2.2 × 10^−10^, **Figure 3E**). Together, these findings suggest that the transition from a CTCF-only state to an enhancer state is associated with relatively modest sequence changes predicted to influence chromatin accessibility, resulting in substantially broader transcription factor occupancy in the enhancer state.

### Dual E/C elements contain architectural and regulatory motifs that exhibit distinct evolutionary signatures

To identify specific transcription factors associated with the transition from the CTCF-only state to the enhancer state, we compared ChromBPNet attribution scores at experimentally supported transcription factor motif instances (Andrews et al. 2023) within dual E/C elements. Across five of the six cell lines, one or more AP-1 family members exhibited significantly higher ChromBPNet attribution scores in enhancer-state than CTCF-only dual E/C elements (*p* < 0.05; **Fig. 4A,B; Supplemental Table S4A**). The specific AP-1 family member exhibiting the strongest enrichment varied among cell lines (JUN, JUND, FOSL2, and/or FOS), likely reflecting both biological and technical variation. HCT116 was the only exception, with similar AP-1 attribution scores in enhancer and CTCF-only states. Additional transcription factors exhibited significant cell type-specific increases in attribution scores, including CEBPB, ATF1, RXRA, FOXA2, and HNF4A (*p* < 0.05). Clustering transcription factors based on overlap among their associated dual E/C elements revealed that, aside from small family clusters such as AP-1 factors, most enriched transcription factors formed distinct groups with limited overlap (**Supplemental Fig. S8**), suggesting that enhancer-state transitions can be achieved through recruitment of different context-specific transcription factors.

To illustrate this process, we examined individual dual E/C elements. For example, EH38E2226676 participates in candidate architectural interactions with the promoter of *PHLDB2* in four of the six cell lines. *PHLDB2* encodes LL5β, a cytoskeletal adaptor that regulates focal adhesion dynamics and cell migration (Lansbergen et al. 2006; Stehbens et al. 2014). In HCT116, EH38E2226676 is in an enhancer state, whereas in the other five cell lines it is in a CTCF-only state (**Fig. 4C, Supplemental Fig. S9**). Across all cell lines, the element exhibits comparable CTCF ChIP-seq and DNase-seq signals accompanied by strong ChromBPNet attribution scores at the CTCF motif. In contrast, HCT116 displays elevated H3K27ac signal together with a notable increase in attribution scores corresponding to a JUN motif, consistent with acquisition of the enhancer state.

A second example is EH38E1475832, a dual E/C element highlighted previously in **Figure 2F** (**Supplemental Fig. S10**). This cCRE is linked through candidate architectural interactions in five of the six cell lines to the promoter of *TYSND1*, which encodes a peroxisomal protease that processes matrix proteins required for peroxisome function and biogenesis (Kurochkin et al. 2007; Mizuno et al. 2013). EH38E1475832 similarly exhibits strong CTCF ChIP-seq and DNase-seq signals across all six cell lines accompanied by high ChromBPNet attribution scores at the CTCF motif. However, enhancer-state classifications in HepG2, MCF-7, and MCF 10A due to elevated H3K27ac signal, coincide with increased ChromBPNet attribution scores at a CEBPB motif.

Our genome-wide analyses and the examples above suggest that enhancer-state dual E/C elements generally comprise two distinct sequence components contributing to accessibility: a recurrent architectural CTCF motif and a more variable enhancer-associated transcription factor motif. We therefore asked whether these architectural and regulatory components exhibit similar evolutionary constraint, as measured using 240-way mammalian PhyloP conservation scores generated by the Zoonomia Consortium (Christmas et al. 2023). Across dual E/C elements, CTCF motif instances had significantly higher PhyloP scores compared with JUN, CEBPB, or other transcription factor motif instances (CTCF median = 2.2 vs. other transcription factor medians −0.24 to 0.28, Wilcoxon test, *p* < 5.9 × 10^−14^; **Fig. 4D**, **Supplemental Fig. S11A**). We observed a similar pattern when we analyzed motif retention across the 240 mammalian genomes, requiring at least 75% sequence identity between orthologous motif instances. Across all paired comparisons, CTCF motifs were consistently retained in more mammalian species than enhancer-associated transcription factor motifs present within the same dual E/C element (mean median difference = 57 species, paired Wilcoxon tests, *p* < 9.5 × 10^−5^; **Fig. 4E, Supplemental Table S4B**).

To illustrate the distinct evolutionary constraints acting on the architectural and enhancer-associated components of dual E/C elements, we revisited EH38E2226676, comparing the conservation of its CTCF and JUN motif sites. The CTCF motif was retained in 197 of the genomes examined, notably only missing in whales and even-toed ungulates (**Fig. 4F, Supplemental Table S4C**). In contrast, the JUN motif was largely restricted to primates, including New World monkeys, Old World monkeys, and great apes (N = 29, **Fig. 4F**). Notably, every species retaining the JUN motif also retained the corresponding CTCF motif. Similar results were observed for the second example EH38E1475832, where the CTCF motif was retained in 229 mammalian genomes compared to 104 genomes retaining the associated CEBPB motif; all species retaining the CEBPB motif also retained the corresponding CTCF motif (**Supplemental Table S4D**). Collectively, these observations support a model in which stable candidate architectural interactions provide evolutionarily conserved regulatory scaffolds that can be repeatedly repurposed across cellular contexts through recruitment of distinct context-specific transcription factors.

### Architectural promoter interactions preferentially connect context-dependent regulatory programs

Across the six cell lines examined in this study, we captured numerous examples of cCREs in candidate architectural interactions transitioning between enhancer and CTCF-only states. However, on average, 81% of cCREs in candidate architectural interactions, which we will refer to as architectural elements, were observed only in the CTCF-only state (**Supplemental Table S5A**). This raised the possibility these architectural elements may acquire enhancer signatures in biological contexts not represented in our current study design. Because the dual E/C analysis above required stringent overlap criteria to confidently identify regulatory state transitions, we next expanded our analysis from the subset of dual E/C elements to all architectural elements.

To characterize the broader enhancer signatures of architectural elements, we quantified H3K27ac enrichment across 562 cell and tissue types in the Registry of cCREs. Overall, 84% of architectural elements exhibited high H3K27ac signal in at least one biological context, indicating that the majority are observed in an enhancer-associated state in at least one surveyed biosample. Because the majority of architectural elements exhibited enhancer-associated states in at least one surveyed biological context, we refer to this broader class as architectural E/C elements throughout the remainder of this section.

Most samples exhibited modest positive enrichment for H3K27ac signals at architectural E/C elements (median = 1.15 fold, **Supplemental Table S5B**), suggesting that context-dependent enhancer signatures at these cCREs are widespread across diverse cellular states. However, we observed notable examples of both enrichment and depletion. Consistent with the enrichment for architectural P-E interactions in HCT116 cells, architectural E/C elements identified in the other five cell lines exhibited consistent enrichment for H3K27ac signal in HCT116-derived biosamples (1.5 - 2.6 fold, average 2.2 fold, **Fig. 5A**). Additionally, architectural E/C elements across all six cell lines were enriched for H3K27ac signal in brain tissue samples (1.6 - 2.4, average 2.1 fold). In contrast, architectural E/C elements were depleted for H3K27ac signals in pluripotent stem cells, developmental progenitor populations, in vitro-derived cellular models, and several immune cell populations, particularly naive and regulatory lymphocyte states (**Fig. 5A**).

**Figure 5.**
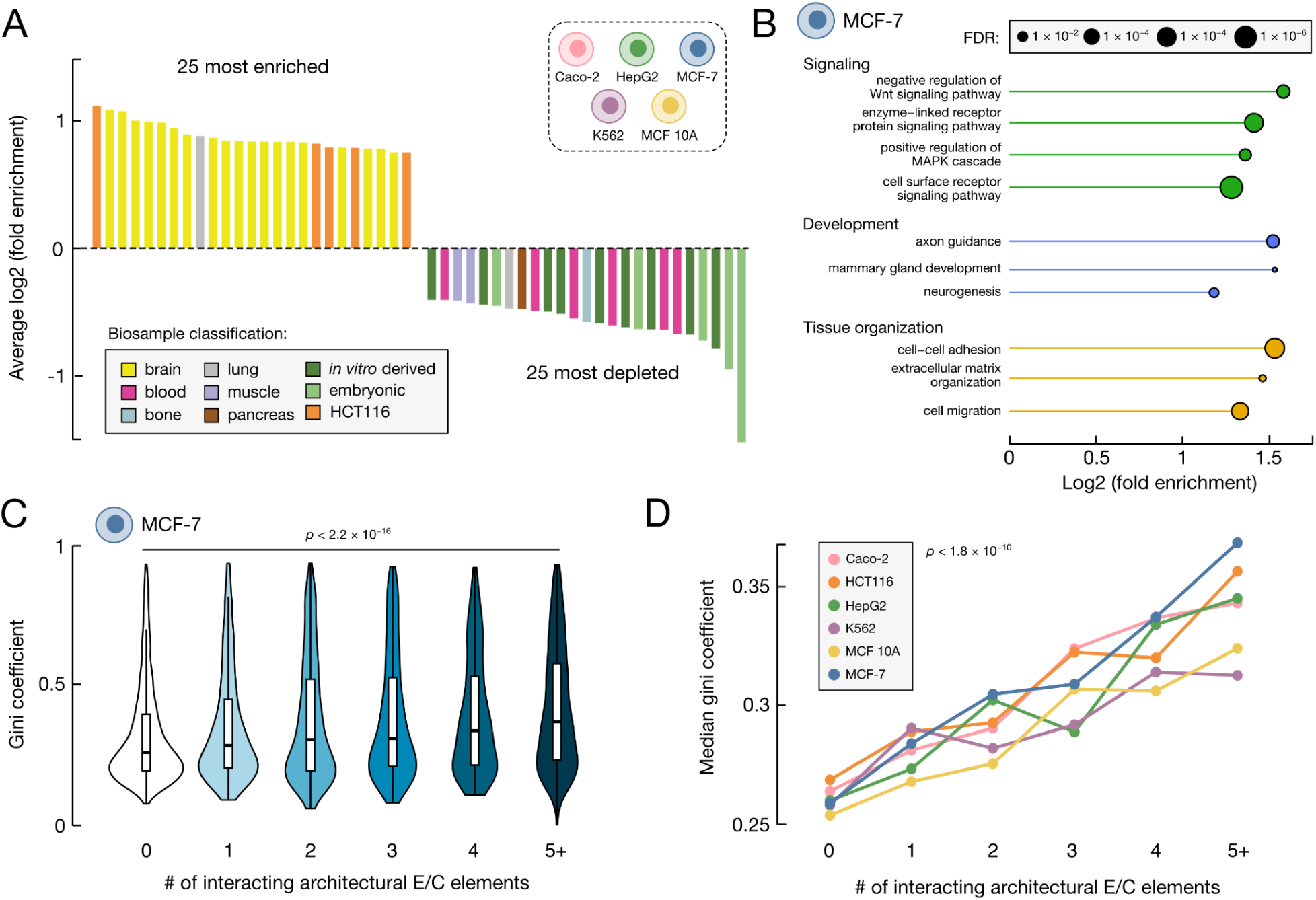
Architectural E/C elements preferentially regulate context-dependent biological programs. (*A*) H3K27ac enrichment for architectural E/C elements across ENCODE biosamples across five cell types (Caco-2, HepG2, K562, MCF 10A, MCF-7). Bars show the 25 most enriched and 25 most depleted biosamples ranked by average log_2_ fold enrichment. Colors indicate biosample classifications, including organ/tissue of origin and developmental state. Corresponding values for all 562 biosamples in each of the six cell lines are provided in **Supplemental Table S5B**. (*B*) Lollipop plots depicting representative Gene Ontology (GO) overrepresentation analysis for genes connected to architectural E/C elements in MCF-7 cells. Terms are grouped by broad categories (signaling: green, development: blue, tissue organization: orange) and circle size indicates false discovery rate (FDR). (*C*) Violin-boxplots depicting the Gini coefficient of genes with promoter-centric interactions in MCF-7 stratified by the number of interacting architectural E/C elements. P-value corresponds to a Jonckheere-Terpstra trend test. Corresponding plots for the other five cell lines are provided in **Supplemental Figure S12**. (*D*) Median Gini coefficients across all six cell lines stratified by the number of interacting architectural E/C elements. Highest p-value from Jonckheere-Terpstra trend tests in each cell line is shown.

To gain insight into the biological programs associated with architectural E/C elements, we performed gene ontology overrepresentation analysis on their linked genes, using genes with any promoter-centric interaction as the background. Across all six cell lines, genes linked to architectural E/C elements were recurrently enriched for developmental regulation, receptor-mediated signaling, cell migration, cell adhesion, and extracellular matrix organization (**Fig. 5B**, **Supplemental Table S5C**). For example in MCF-7 cells, enriched terms included cell-cell adhesion, cell migration, receptor signaling, Wnt/MAPK signaling, mammary gland development, axon guidance, and neurogenesis (**Fig. 5B**).

These enriched gene ontology terms suggested that architectural E/C elements preferentially connect genes with context-dependent expression patterns. To test this hypothesis, we quantified expression specificity using Gini coefficients calculated from Bru-seq data across 16 deeply profiled cell lines (Moore et al. 2026). Promoters connected to at least one architectural E/C element exhibited higher Gini coefficients than promoters connected to other cCRE classes, consistent with more specific expression across cell types (0.30 vs 0.26, Wilcoxon test, *p* < 2.0 × 10^−9^). Expression specificity also increased progressively with the number of interacting architectural E/C elements (Jonckheere-Terpstra trend test, p < 2.2 × 10^−16^; **Fig. 5C,D, Supplemental Fig. S12**). This relationship was reproducible in each individual cell line, with promoters connected to five or more architectural E/C elements consistently exhibiting the highest Gini coefficients (average of 0.34).

Together, these observations suggest that architectural E/C elements represent a broadly used regulatory strategy that is preferentially leveraged in specific biological contexts. Across diverse cell and tissue types, these elements interact with genes involved in signaling, development, and other context-dependent biological programs, with increasing architectural connectivity associated with increasingly specialized patterns of gene expression. More broadly, these results illustrate how integrating chromatin interaction maps with regulatory annotations and architectural organization can reveal biologically meaningful classes of chromatin interactions that would be difficult to recognize using any single data type alone.

## Discussion

Three-dimensional chromatin interaction maps have become increasingly comprehensive, yet understanding what individual chromatin interactions represent biologically remains a major challenge. By integrating complementary genomic modalities, we found that promoter-centric chromatin interactions comprise multiple categories with distinct structural and regulatory properties. Candidate architectural interactions emerged as a particularly distinctive class, exhibiting increased recurrence across cellular contexts and context-dependent regulatory activity. Notably, many cCREs in these candidate architectural interactions transitioned between CTCF-only and enhancer states while maintaining stable chromatin connections, demonstrating that architectural and enhancer states can frequently coexist within the same regulatory element. Collectively, these findings suggest that chromatin architecture provides a stable framework for context-dependent regulatory activity rather than serving solely as a static structural feature of genome organization.

Our evolutionary analyses suggest a possible explanation for how new regulatory functions arise at existing architectural interactions. Within dual E/C elements, CTCF motifs were much more conserved than enhancer-associated transcription factor motifs, both in their sequence conservation and in the number of mammalian species retaining the motif. These observations are consistent with the hypothesis that architectural interactions may provide stable regulatory scaffolds onto which new transcription factor binding sites can accumulate over evolutionary time while maintaining their underlying chromatin interactions. Although present-day genomes cannot establish the evolutionary order of these events, our findings are consistent with this model. Testing this hypothesis will require additional comparative genomics together with experimental studies of recently evolved regulatory elements.

Our findings also suggest a possible explanation for why some enhancers utilize architectural interactions. Compared with their non-architectural counterparts, architectural interactions span longer genomic distances, are more recurrent across cell types, and more often connect enhancers to distal rather than nearby genes. Together, these properties suggest that architectural interactions provide stable, long-range connections that can be repeatedly reused across diverse cellular contexts. Such pre-existing interactions may be particularly advantageous for genes involved in signaling, development, or other context-dependent processes by increasing the likelihood that regulatory elements and their target promoters are already in close spatial proximity when additional regulatory inputs are acquired. Context-dependent transitions of architectural elements to the enhancer state may therefore explain how highly recurrent physical interactions give rise to more specific patterns of gene regulation.

These advantages may, however, come at the cost of reduced regulatory specificity. Current models of cohesin-mediated loop extrusion explain how stable chromatin loops are established through CTCF binding and motif orientation. However, both our analyses and previous studies indicate that, across cell populations, individual CTCF-bound elements are observed interacting with multiple convergently oriented CTCF sites, suggesting that loop extrusion naturally generates multiple potential long-range contacts rather than a single fixed interaction. Consistent with this model, enhancers participating in candidate architectural interactions interacted with more promoters on average than non-architectural enhancers. Together, these observations suggest that architectural interactions may define a broader set of potential regulatory partners rather than a single target gene. If so, additional mechanisms—including promoter-enhancer compatibility (Zabidi et al. 2015; Martinez-Ara et al. 2022; Bergman et al. 2022), promoter responsiveness (Tan et al. 2026), cell type-specific transcription factor recruitment, or other regulatory constraints—are likely required to determine which of these physical interactions ultimately functions as a regulatory interaction.

Our analyses suggest that architectural interactions are broadly leveraged for distal gene regulation across diverse cellular programs. We identified dual E/C elements in each of the six cell lines examined and extended this analysis to define architectural E/C elements across hundreds of additional cell and tissue types represented in the Registry of cCREs. At the same time, the enrichment and depletion patterns observed across biosamples suggest that different biological systems rely on this regulatory strategy to different extents. For example, architectural E/C elements were consistently depleted in pluripotent stem cells, developmental progenitor populations, and multiple immune cell types, particularly those in naive or unstimulated states. One explanation for this consistent depletion is that architectural E/C elements are more likely to mediate transitions between cellular states, whereas enhancers maintaining stable pluripotent or naive states may rely more heavily on non-architectural contacts. In this model, architectural elements would remain largely in a CTCF-only state until developmental or environmental signals activate enhancer programs that promote a new regulatory state. This model is also consistent with our observation that genes linked to architectural E/C elements were enriched for signaling and developmental functions and exhibited increased expression specificity across cell types. Testing this model will require differentiation and stimulation time courses that track architectural interactions, enhancer activity, and transcriptional responses within the same system.

Brain tissue samples provided the clearest example of the opposite pattern, exhibiting consistent enrichment for enhancer signatures at architectural E/C elements. Although the enrichment observed in bulk brain tissue likely reflects contributions from multiple brain cell types, it was also accompanied by enrichment of neural developmental and signaling functions among linked genes, together suggesting that neuronal regulatory programs may preferentially leverage architectural interactions for gene regulation. Neuronal genes are often exceptionally long (Zylka et al. 2015) and require precise regulation across diverse cell types and developmental stages, properties that may particularly benefit from stable architectural interactions capable of supporting long-range, context-dependent regulation. If so, disruption of chromatin architecture would be expected to disproportionately affect nervous system development and function and could explain some of the neurological phenotypes associated with cohesinopathies (Piché et al. 2019) and CTCF-related disorders (Valverde de Morales et al. 1993, 2023). Future studies integrating chromatin architecture, enhancer activity, and perturbation experiments in neuronal systems will be important for determining how architectural E/C elements contribute to nervous system development and disease.

The organizational differences between architectural and non-architectural interactions also have important implications for computational models of enhancer-gene regulation. Consistent with previous studies, assigning enhancers to their nearest gene correctly identifies many regulatory relationships but fails to recover a substantial fraction of distal interactions, particularly those mediated by architectural interactions. This likely explains why computational methods that integrate chromatin interaction information with epigenomic measurements, such as ABC (Fulco et al. 2019) and rE2G (Gschwind et al. 2023), achieve improved performance in predicting functional enhancer-promoter relationships. Our results also have implications for the development and evaluation of future computational methods. Because architectural and non-architectural interactions differ in their genomic distances, recurrence across cell types, and target-gene selection, different computational approaches may perform differently across these interaction classes. Benchmarking efforts, such as our own BENGI resource (Moore et al. 2020), should therefore consider architectural organization alongside traditional performance metrics to better understand the strengths and limitations of enhancer-gene prediction methods in different regulatory contexts. More broadly, incorporating biologically meaningful interaction subclasses into computational models may improve both enhancer-gene prediction and our understanding of how different regulatory mechanisms contribute to gene regulation.

Although our framework provides new insight into chromatin interaction organization, several important questions remain. First, our analyses were performed across six deeply characterized human cell lines, five of which are transformed cell lines with abnormal karyotypes. While these models do not capture the full spectrum of normal human biology, they provide an unusually rich collection of complementary functional genomic datasets that enabled systematic integration of chromatin interactions, regulatory annotations, chromatin accessibility, transcription factor binding, and transcriptional activity. Extending this framework to additional primary cell types, tissues, and developmental systems will be important for determining how broadly these patterns generalize. Second, chromatin interaction datasets remain subject to substantial technical variability arising from differences in assay sensitivity, sequencing depth, and interaction-calling approaches. Consequently, the absence of a detected interaction in one cell type should not necessarily be interpreted as evidence that the interaction is absent biologically. Throughout this study, we sought to minimize these effects by emphasizing comparisons within individual cell types and by focusing analyses on recurrent interaction patterns observed across multiple biological contexts. Continued improvements in chromatin interaction assays and increasingly comprehensive interaction maps should further refine this framework. Finally, the chromatin interaction assays used throughout this study measure physical proximity rather than direct regulatory function. Although chromatin interactions provide a powerful proxy for identifying candidate regulatory relationships, not every physical interaction is expected to contribute to gene regulation. Future studies integrating functional perturbation assays, such as CRISPR-based enhancer perturbations, with chromatin interaction maps will help determine which predicted interactions have measurable regulatory consequences. At the same time, functional perturbation assays introduce their own limitations, including enhancer redundancy and context-dependent effects. Rather than replacing chromatin interaction data, these complementary approaches should provide a more complete understanding of how three-dimensional genome organization contributes to gene regulation.

Over the past two decades, functional genomics has transformed our ability to identify and annotate individual cis-regulatory elements across the human genome. A remaining challenge is understanding how these elements function together to regulate gene expression. Our results suggest that meeting this challenge will require not only increasingly complete chromatin interaction maps but also biologically informed frameworks for interpreting the interactions between cis-regulatory elements. Just as cis-regulatory element annotations provide insight into the function of individual genomic loci, interaction annotations provide insight into how regulatory elements work together to control transcription. Such frameworks will provide a richer understanding of how genome organization contributes to gene regulation in development, cellular identity, and human disease.

## Methods

### Data acquisition

The sources and accessions for datasets analyzed in this study are listed in **Supplemental Table S1A**. Cell type specific cCREs for each of the six cells lines were downloaded from SCREEN (V4 of the Registry, 2026, screen.wenglab.org) and Hi-C loop calls, RNA-seq and Bru-seq quantifications, and ChromBPNet attribution models were downloaded directly from the ENCODE Portal (encodeproject.org). RNAPII and CTCF ChIA-PET interactions were downloaded from the ENCODE Portal and processed using the standardized filtering strategy described in (Moore et al. 2026). These filtered ChIA-PET interaction annotations, together with all derived annotations generated in this study, are available at data.moore-lab.org.

### Comparison of chromatin interaction assays

Within each cell line, we calculated pairwise overlaps among the three-dimensional chromatin assays, and interactions were classified according to the combination of assays in which they were detected (RNAPII ChIA-PET only, CTCF ChIA-PET only, Hi-C only, RNAPII ChIA-PET & CTCF ChIA-PET, RNAPII ChIA-PET & Hi-C, CTCF ChIA-PET & Hi-C, or all three assays). To calculate overlap we used bedtools (v2.30.0; (Quinlan and Hall 2010), requiring both interaction anchors to overlap between datasets. Interactions unique to a given assay were defined as those that did not overlap an interaction detected by either of the other two assays. Counts and proportions for each overlap category were summarized separately for each assay and cell line.

### Extension of cCRE classifications

The Registry of cCREs classifies regulatory elements using combinations of chromatin accessibility, histone modification, and CTCF occupancy. Because many CTCF-bound elements exhibit relatively low chromatin accessibility, we extended this classification scheme by introducing an additional CTCF-bound cCRE class that does not require high DNase-seq signal. Specifically, for each biosample, cCREs with a normalized CTCF ChIP-seq signal z-score greater than 1.64 (one-sided 95th percentile) were designated as CTCF-bound cCREs, irrespective of their chromatin accessibility or histone modification status. We also designated high signal CTCF-bound cCREs (for the purposes of identifying architectural interactions) as those with signal z-scores greater than 2.32 (one-sided 99th percentile). These additional categories were used throughout the present study to identify CTCF-associated regulatory elements and candidate architectural interactions, respectively, while preserving the original Registry classifications for all other cCRE classes.

### Identification and validation of candidate architectural interactions

To characterize the relationship between chromatin interactions and architectural protein binding, we quantified CTCF occupancy at interaction anchors across all Hi-C, RNAPII ChIA-PET, and CTCF ChIA-PET datasets. We intersected chromatin interactions with high signal CTCF-bound cCREs (z-score > 2.32) using bedtools, requiring both interacting anchors to overlap at least one high signal CTCF-bound cCRE. To account for modest differences in interaction anchor placement among assays, interactions with both anchors overlapping a candidate architectural interaction identified in the previous step were also assigned candidate architectural support. All remaining interactions were classified as non-architectural.

To evaluate this approach, we compared it with alternative definitions based on cohesin occupancy and CTCF motif orientation (**Supplemental Methods**). Cohesin-supported interactions were defined as interactions in which both anchors overlapped RAD21 ChIP-seq peaks. Convergently oriented interactions were identified by intersecting interaction anchors with forward and reverse CTCF motif instances from representative transcription factor peaks (**Supplemental Dataset S4**) and identifying interactions in which the two anchors contained motifs in convergent orientation. We quantified agreement amongst the different approaches by comparing the overlap of interactions classified by each (**Supplemental Table S1E**).

### Identification and regulatory classification of promoter-centric interactions

To classify promoter-centric chromatin interactions, Hi-C, RNAPII ChIA-PET, and CTCF ChIA-PET interactions were aggregated within each cell line and intersected with cell type-specific cCRE annotations using bedtools. All interactions with at least one promoter cCRE were retained for analysis and classified according to the following hierarchy:

- Promoter–promoter (P–P): both interaction anchors overlapped promoter cCREs.
- Promoter–enhancer (P–E): one anchor overlapped a promoter cCRE and the second anchor overlapped at least one proximal and/or distal enhancer cCRE.
- Promoter–CTCF (P–C): one anchor overlapped a promoter cCRE and the second anchor overlapped a CTCF-bound cCRE.
- Promoter–other (P–O): one anchor overlapped a promoter cCRE and the second anchor overlapped another active non-promoter, non-CTCF-bound cCRE (e.g. CA-H3K4me3, CA-TF, CA, TF).
- Promoter–inactive (P–I): one anchor overlapped a promoter cCRE and the second anchor overlapped an inactive (low chromatin accessibility) cCRE.
- Promoter–none (P–N): one anchor overlapped a promoter cCRE and the second anchor did not overlap any annotated cCRE.

Categories were assigned hierarchically, with promoter and enhancer annotations taking precedence over CTCF and other cCRE annotations, thereby ensuring that each interaction was assigned to a single, non-overlapping promoter-centric class.

### Comparison of interaction distances

For each promoter-centric interaction, we calculated its genomic span as the distance between the end coordinate of the first interaction anchor and the start coordinate of the second interaction anchor. We compared distance distributions separately for P-P, P-E, and P-C interactions after stratifying interactions as candidate architectural or non-architectural using two-sided Wilcoxon rank-sum tests.

### Interaction recurrence across cell lines

For each P–P, P–E, and P–C interaction in a given cell line, we calculated its recurrence (i.e., the number of cell lines in which the interaction was detected), using bedtools. We considered an interaction recurrent if both of its anchors overlapped a promoter-centric interaction in another cell line, regardless of its regulatory classification. Recurrence was summarized separately for each promoter-centric interaction class by calculating the fraction of interactions detected in at least four of the six cell lines.

### Gene expression associated with promoter-centric interactions

We assigned genes with active promoters to three groups in each cell line: genes whose promoters participated in at least one P–P interaction, genes whose promoters participated exclusively in P–E and/or P–C interactions, and genes with active promoters that did not participate in any P–P, P–E, or P–C interaction. We then quantified the expression of genes in each group using total RNA-seq TPM values, averaged across biological replicates for each cell line. Finally, we compared expression distributions among groups using pairwise Wilcoxon rank-sum tests with FDR correction.

### Promoter-promoter co-expression

To evaluate co-expression between genes connected by promoter-promoter interactions, we calculated Pearson correlation coefficients using log-transformed Bru-seq RPKM values across 16 deeply profiled cell lines. As a background comparison, we generated randomly selected pairs of genes with active promoters in the corresponding cell line. Correlation distributions for observed P–P pairs and random active-promoter pairs were compared using two-sided Wilcoxon rank-sum tests.

### Evaluation of closest-gene assignments

For each enhancer or CTCF-bound cCRE, we determined whether the interacting promoter corresponded to its closest gene using multiple commonly used closest-gene definitions and transcript annotations (detailed in the **Supplemental Methods**). cCREs not connected to their closest gene were further classified according to the distance to the nearest TSS. Those located within 10 kb of the nearest TSS were designated unresolved, as chromatin interactions at these distances cannot be reliably detected. The remaining cCREs were classified as interacting exclusively with distal genes.

### Benchmarking closest-expressed-gene prediction

To evaluate a simple proximity-based assignment strategy, we implemented a closest-expressed-gene heuristic for distal enhancer and CTCF-bound cCREs participating in promoter-centric interactions. For each cell line, we quantified gene expression using total RNA-seq TPM values averaged across biological replicates and defined expressed genes as genes with TPM > 1. For each cCRE, we assigned the predicted target gene as the expressed gene with the nearest TSS based on linear proximity. We then compared this closest-expressed-gene prediction with the promoter connected by the observed chromatin interaction. We summarized performance as the fraction of cCREs for which the closest expressed gene matched an observed interacting promoter and the fraction of observed promoter-centric interactions recovered by the closest-expressed-gene method. cCREs located within 10 kb of the nearest TSS were excluded from this benchmark because interactions at these distances fall below the reliable resolution of current chromatin interaction assays.

### Regulatory category switching across cell lines

We selected candidate architectural P-E interactions in each cell line and compared them with promoter-interactions across the remaining cell lines using bedtools. Interactions were considered equivalent when both anchors overlapped. For each matched interaction, we recorded the regulatory category of the interaction (P-E, P-C, or other) in the comparison cell line. This analysis quantified the frequency with which candidate architectural promoter-enhancer interactions retained enhancer annotations or transitioned to CTCF-bound states across cellular contexts.

### Definition of dual E/C elements

To distinguish whether transitions between P–E and P–C annotations reflected adjacent regulatory elements or state changes within the same cCRE, we first identified all candidate architectural P–E interactions that were also classified as P–C interactions in another cell line (as described above). We then restricted this set to interactions whose enhancer anchor overlapped a single enhancer cCRE, yielding 1,499 unique enhancer cCREs. Finally, we determined whether these enhancer cCREs corresponded to the same CTCF-bound cCRE in the matched P–C interactions across the other cell lines. Enhancer cCREs satisfying this criterion were designated dual E/C elements (n = 946).

### CAPRA enhancer activity quantification

To evaluate enhancer activity, we used previously generated per-cCRE CAPRA STARR-seq scores from K562 cells (Moore et al. 2026). Dual E/C elements were stratified according to their state in K562: enhancer vs. CTCF-only. We then extracted CAPRA scores for enhancer-state dual E/C elements, CTCF-only dual E/C elements, all K562 enhancer cCREs, and a randomly selected background set of cCREs. Score distributions were compared using two-sided pairwise Wilcoxon rank-sum tests with FDR correction.

### ChromBPNet high-attribution base quantification

To compare sequence-level regulatory contributions between enhancer and CTCF-only states of dual E/C elements, we analyzed cell type-specific ChromBPNet attribution scores across cCREs. For each dual E/C element, we identified bases with high attribution scores, defined as bases with attribution values greater than or equal to 10% of the maximum attribution score within that cCRE. We then calculated both the total number of high-attribution bases and the fraction of each cCRE comprising high-attribution bases. Dual E/C elements were stratified according to their regulatory state in each cell line (e.g. enhancer-state or CTCF-only-state), Distributions were compared between states using two-sided Wilcoxon rank-sum tests.

### Transcription factor ChIP-seq summit overlap

To compare transcription factor occupancy between enhancer and CTCF-only states of dual E/C elements, we intersected dual E/C elements with ENCODE transcription factor ChIP-seq peak summits in HepG2 and K562 cells (**Supplemental Data S3**). For each dual E/C element, we counted the number of overlapping transcription factor ChIP-seq peak summits using bedtools. Distributions of summit counts were compared between enhancer-state and CTCF-only-state dual E/C elements using two-sided Wilcoxon rank-sum tests.

### ChromBPNet motif attribution analysis

To identify transcription factors associated with enhancer-state transitions, we quantified ChromBPNet attribution scores at experimentally supported transcription factor motif instances within dual E/C elements. Using bedtools, we intersected representative transcription factor motif instances (**Supplemental Data S4**) with dual E/C elements, restricting the analysis to transcription factors represented by at least 100 experimentally supported motif instances. For each cell line, we then calculated the average ChromBPNet attribution score across each motif instance using UCSC tools bigWigAverageOverBed (Kuhn et al. 2013) and cell type-specific ChromBPNet attribution bigWig files. Stratifying motif instances by enhancer and CTCF-only states, we evaluated differences in attribution scores using two-sided Wilcoxon rank-sum tests. Transcription factors exhibiting significant differences (*p* < 0.05) in at least one cell line were retained for downstream analyses and visualization.

### Evolutionary conservation of transcription factor motifs

We compared the evolutionary conservation of transcription factor motifs at dual E/C elements using two approaches. First, we compared base pair-level evolutionary constraint using 240-mammal PhyloP scores generated by the Zoonomia Project (Christmas et al. 2023). For experimentally validated motif instances that overlapped dual E/C elements (as identified above) we calculated (i) average PhyloP scores across each motif using bigWigAverageOverBed and (ii) generated aggregate PhyloP profiles centered on motif instances using deepTools computeMatrix (Ramírez et al. 2014).

Second, to compare motif retention across species, we compared CTCF vs. other transcription factor motif instances occurring within the same dual E/C element. We then queried the Zoonomia HDF5 multiple genome alignment (Andrews et al. 2023) to determine, for each motif instance, the number of mammalian species in which the orthologous motif sequence retained at least 75% sequence identity relative to the human reference sequence. Paired comparisons between CTCF and enhancer-associated transcription factor motifs within the same dual E/C element were evaluated using two-sided paired Wilcoxon signed-rank tests. Summary statistics were calculated separately for each transcription factor and reported as median numbers of retained species.

### H3K27ac enrichment at architectural E/C elements across Registry biosamples

To assess enhancer-associated activity across a broad range of biological contexts, we quantified H3K27ac enrichment at architectural E/C elements using the expanded Registry of cCREs. For each cell line, architectural E/C elements were defined as CTCF-bound cCREs participating in candidate architectural P-C interactions. H3K27ac enrichment was calculated across Registry biosamples by comparing the fraction of architectural E/C elements with high H3K27ac signal to the corresponding fraction across all cCREs. Fold enrichment was calculated as:

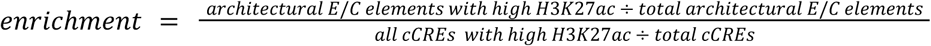

Enrichment values were calculated separately for architectural E/C elements defined in each of the six cell lines and summarized across 525 biosamples.

### Gene ontology enrichment

To identify biological programs associated with architectural E/C elements, we performed Gene Ontology enrichment analysis using PANTHER (v19.0 (Thomas et al. 2022; Mi et al. 2019)). For each cell line, the foreground gene set consisted of genes whose promoters were connected to architectural E/C elements through candidate architectural P–E or P–C interactions. Cell line-specific background sets consisted of all genes whose promoters participated in any P–P, P–E, or P–C interaction in that cell line. Gene identifiers were collapsed to gene-level Ensembl IDs before enrichment analysis. Foreground and background gene lists were submitted to PANTHER separately for each cell line using default parameters. Enrichment analyses were performed across all three Gene Ontology domains: Biological Process, Molecular Function, and Cellular Component.

### Expression specificity analysis

To evaluate whether architectural E/C elements preferentially connect genes with context-dependent expression patterns, we analyzed gene-level Gini coefficients calculated from quantile-normalized Bru-seq expression measurements across 16 deeply profiled cell lines (Moore et al. 2026). For each of the six cell lines with interaction data, genes were stratified by the number of interacting architectural E/C elements connected to their promoters through candidate architectural P–E or P–C interactions. Genes with P–E or P–C interactions but no architectural E/C elements were assigned to the zero-interaction group. Gini coefficient distributions were compared across groups containing 0, 1, 2, 3, 4, or ≥5 interacting architectural E/C elements. Monotonic trends between the number of interacting architectural E/C elements and expression specificity were evaluated using Jonckheere-Terpstra trend tests.

## Data availability

All data and annotations generated by this study are available as Supplemental Datasets and available to download at https://data.moore-lab.org. Annotations are also available as a track hub for the UCSC genome browser (https://users.moore-lab.org/Track-Hubs/3D-Interactions/hub.txt) and can also be visualized at https://hubble.moore-lab.org.

## Code availability

All code used in this study available at https://github.com/Moore-Lab-UMass/3D-Interactions

## Acknowledgements

This study was supported by National Institutes of Health grants R03OD036496, R35GM160300 U24HG009446 and U24HG012343.

We thank the ENCODE Registry of cCRE Working Group for providing valuable, early-stage feedback on this work.

## Ethics declarations (competing interests)

A.K. is a consulting fellow with Illumina; a member of the SABs of OpenTargets (GSK), PatchBio, and SerImmune

## Author contributions

J.E.M. conceived the study, designed and performed computational analyses, interpreted the results, and wrote the manuscript. M.C. performed computational analyses, prepared figures, and contributed to manuscript writing. G.A. and M.G. contributed annotation resources, including transcription factor motif and comparative genomics datasets, used in the downstream analyses. A.K. contributed to conceptual discussions, interpretation of the ChromBPNet analyses, and manuscript editing. V.K. and V.A. contributed to the development, training, and dissemination of the ChromBPNet models used in this study. All authors reviewed and approved the final manuscript.

## References

Andrews G, Fan K, Pratt HE, Phalke N, Zoonomia Consortium§, Karlsson EK, Lindblad-Toh K, Gazal S, Moore JE, Weng Z. 2023. Mammalian evolution of human cis-regulatory elements and transcription factor binding sites. Science 380: eabn7930.

Bergman DT, Jones TR, Liu V, Ray J, Jagoda E, Siraj L, Kang HY, Nasser J, Kane M, Rios A, et al. 2022. Compatibility rules of human enhancer and promoter sequences. Nature 607: 176–184.

Christmas MJ, Kaplow IM, Genereux DP, Dong MX, Hughes GM, Li X, Sullivan PF, Hindle AG, Andrews G, Armstrong JC, et al. 2023. Evolutionary constraint and innovation across hundreds of placental mammals. Science 380: eabn3943.

Fulco CP, Nasser J, Jones TR, Munson G, Bergman DT, Subramanian V, Grossman SR, Anyoha R, Doughty BR, Patwardhan TA, et al. 2019. Activity-by-contact model of enhancer-promoter regulation from thousands of CRISPR perturbations. Nat Genet 51: 1664–1669.

Fullwood MJ, Liu MH, Pan YF, Liu J, Xu H, Mohamed YB, Orlov YL, Velkov S, Ho A, Mei PH, et al. 2009. An oestrogen-receptor-alpha-bound human chromatin interactome. Nature 462: 58–64.

Gschwind AR, Mualim KS, Karbalayghareh A, Sheth MU, Dey KK, Jagoda E, Nurtdinov RN, Xi W, Tan AS, Jones H, et al. 2023. An encyclopedia of enhancer-gene regulatory interactions in the human genome. Genomics. https://www.biorxiv.org/content/10.1101/2023.11.09.563812v1.

Kuhn RM, Haussler D, Kent WJ. 2013. The UCSC genome browser and associated tools. Brief Bioinform 14: 144–161.

Kurochkin IV, Mizuno Y, Konagaya A, Sakaki Y, Schönbach C, Okazaki Y. 2007. Novel peroxisomal protease Tysnd1 processes PTS1- and PTS2-containing enzymes involved in beta-oxidation of fatty acids. EMBO J 26: 835–845.

Lansbergen G, Grigoriev I, Mimori-Kiyosue Y, Ohtsuka T, Higa S, Kitajima I, Demmers J, Galjart N, Houtsmuller AB, Grosveld F, et al. 2006. CLASPs attach microtubule plus ends to the cell cortex through a complex with LL5beta. Dev Cell 11: 21–32.

Lieberman-Aiden E, van Berkum NL, Williams L, Imakaev M, Ragoczy T, Telling A, Amit I, Lajoie BR, Sabo PJ, Dorschner MO, et al. 2009. Comprehensive mapping of long-range interactions reveals folding principles of the human genome. Science 326: 289–293.

Li G, Ruan X, Auerbach RK, Sandhu KS, Zheng M, Wang P, Poh HM, Goh Y, Lim J, Zhang J, et al. 2012. Extensive promoter-centered chromatin interactions provide a topological basis for transcription regulation. Cell 148: 84–98.

Martinez-Ara M, Comoglio F, van Arensbergen J, van Steensel B. 2022. Systematic analysis of intrinsic enhancer-promoter compatibility in the mouse genome. Mol Cell 82: 2519–2531.e6.

Mi H, Muruganujan A, Huang X, Ebert D, Mills C, Guo X, Thomas PD. 2019. Protocol Update for large-scale genome and gene function analysis with the PANTHER classification system (v.14.0). Nat Protoc 14: 703–721.

Mizuno Y, Ninomiya Y, Nakachi Y, Iseki M, Iwasa H, Akita M, Tsukui T, Shimozawa N, Ito C, Toshimori K, et al. 2013. Tysnd1 deficiency in mice interferes with the peroxisomal localization of PTS2 enzymes, causing lipid metabolic abnormalities and male infertility. PLoS Genet 9: e1003286.

Moore JE, Pratt HE, Fan K, Phalke N, Fisher J, Elhajjajy SI, Andrews G, Gao M, Shedd N, Fu Y, et al. 2026. An expanded registry of candidate cis-regulatory elements. Nature 1–10.

Moore JE, Pratt HE, Purcaro MJ, Weng Z. 2020. A curated benchmark of enhancer-gene interactions for evaluating enhancer-target gene prediction methods. Genome Biol 21: 17.

Pampari A, Shcherbina A, Kvon EZ, Kosicki M, Nair S, Kundu S, Kathiria AS, Risca VI, Kuningas K, Alasoo K, et al. 2025. ChromBPNet: bias factorized, base-resolution deep learning models of chromatin accessibility reveal cis-regulatory sequence syntax, transcription factor footprints and regulatory variants. bioRxivorg. 10.1101/2024.12.25.630221.

Piché J, Van Vliet PP, Pucéat M, Andelfinger G. 2019. The expanding phenotypes of cohesinopathies: one ring to rule them all! Cell Cycle 18: 2828–2848.

Quinlan AR, Hall IM. 2010. BEDTools: a flexible suite of utilities for comparing genomic features. Bioinformatics 26: 841–842.

Ramírez F, Dündar F, Diehl S, Grüning BA, Manke T. 2014. deepTools: a flexible platform for exploring deep-sequencing data. Nucleic Acids Res 42: W187–91.

Rao SSP, Huntley MH, Durand NC, Stamenova EK, Bochkov ID, Robinson JT, Sanborn AL, Machol I, Omer AD, Lander ES, et al. 2014. A 3D map of the human genome at kilobase resolution reveals principles of chromatin looping. Cell 159: 1665–1680.

Rowley MJ, Corces VG. 2018. Organizational principles of 3D genome architecture. Nat Rev Genet 19: 789–800.

Sanyal A, Lajoie BR, Jain G, Dekker J. 2012. The long-range interaction landscape of gene promoters. Nature 489: 109–113.

Stehbens SJ, Paszek M, Pemble H, Ettinger A, Gierke S, Wittmann T. 2014. CLASPs link focal-adhesion-associated microtubule capture to localized exocytosis and adhesion site turnover. Nat Cell Biol 16: 561–573.

Tang Z, Luo OJ, Li X, Zheng M, Zhu JJ, Szalaj P, Trzaskoma P, Magalska A, Wlodarczyk J, Ruszczycki B, et al. 2015. CTCF-mediated human 3D genome architecture reveals chromatin topology for transcription. Cell 163: 1611–1627.

Tan Y, Ray J, Sheth MU, Doughty BR, Greenleaf WJ, Engreitz JM. 2026. Intrinsic promoter responsiveness dictates sensitivity to transcriptional activation by enhancers. bioRxivorg 2026.06.25.734173. https://www.biorxiv.org/content/10.64898/2026.06.25.734173v1.full.

The ENCODE Project Consortium. 2026. The Encyclopedia of DNA elements. bioRxiv 2026.07.06.731365. https://www.biorxiv.org/content/10.64898/2026.07.06.731365v1.

Thomas PD, Ebert D, Muruganujan A, Mushayahama T, Albou L-P, Mi H. 2022. PANTHER: Making genome-scale phylogenetics accessible to all. Protein Sci 31: 8–22.

Uyehara CM, Apostolou E. 2023. 3D enhancer-promoter interactions and multi-connected hubs: Organizational principles and functional roles. Cell Rep 42: 112068.

Valverde de Morales HG, Wang H-L, Garber K, Corces V, Li H. 1993. CTCF-related disorder. University of Washington, Seattle, Seattle.

Valverde de Morales HG, Wang H-LV, Garber K, Cheng X, Corces VG, Li H. 2023. Expansion of the genotypic and phenotypic spectrum of CTCF-related disorder guides clinical management: 43 new subjects and a comprehensive literature review. Am J Med Genet A 191: 718–729.

Zabidi MA, Arnold CD, Schernhuber K, Pagani M, Rath M, Frank O, Stark A. 2015. Enhancer-core-promoter specificity separates developmental and housekeeping gene regulation. Nature 518: 556–559.

Zylka MJ, Simon JM, Philpot BD. 2015. Gene length matters in neurons. Neuron 86: 353–355.

